# Serial acquisition of virulence determinants and aminoglycoside resistance by the emerging *Streptococcus agalactiae* sequence type 1010 lineage

**DOI:** 10.64898/2026.08.17.745249

**Authors:** Anne Claire Farid, Sydney Haldeman, Caitlin Otto, Adonis D’Mello, Hervé Tettelin, Adam J. Ratner

**Affiliations:** Department of Pediatrics, NYU Grossman School of Medicine, New York, NY, USA; Department of Pathology, NYU Grossman School of Medicine, New York, NY, USA; Department of Microbiology and Immunology, Institute for Genome Sciences, University of Maryland School of Medicine, Baltimore, Maryland, USA; Department of Microbiology, NYU Grossman School of Medicine, New York, NY, USA

**Keywords:** Group B *Streptococcus*, *Streptococcus agalactiae*, epidemiology, antimicrobial resistance

## Abstract

Based on recent epidemiologic studies, *Streptococcus agalactiae* (Group B *Streptococcus*; GBS) sequence type (ST) 1010 is an emerging lineage now identified in multiple countries. We report the phylogenetic and genomic characteristics of a set of 55 GBS sequence type (ST) 1010 strains, as well as two newly described single-locus variants of ST1010. A core genome phylogeny suggests that ST1010 is closely related to both ST452 and the hypervirulent clonal complex (CC) 17 GBS lineage. Notably, we demonstrate that genes encoding two virulence determinants previously described as specific to CC17 GBS, the HvgA adhesin and the serine-rich repeat protein Srr2, are both present in ST1010 genomes. Srr2 is shared with members of ST452. High-level gentamicin resistance (HLGR) encoded on an IS256 mobile element, previously described in a small number of ST1010 isolates, is present in a distinct ST1010 subclade encompassing the majority of ST1010 isolates. The relationship between ST452 (serotype IV), ST1010 (serotype IV), and ST17 (serotype III) strains suggests that ST17 may have arisen from a serotype IV ancestor and later acquired the type III capsule locus. Taken together, these findings clarify the phylogenetic position of ST1010 and suggest sequential acquisition of virulence determinants and HLGR prior to its international emergence.

**IMPACT STATEMENT:** ST1010 GBS has emerged internationally, with colonizing and invasive isolates described in the United States, Dominican Republic, Netherlands, and Italy. Using a core genome phylogeny and targeted detection of genomic regions, we demonstrate that ST1010 shares specific virulence determinants with the CC17 hypervirulent GBS lineage and that HLGR is confined to a specific numerically dominant subclade of ST1010. Our work spotlights the importance of future epidemiologic and genomic surveillance of ST1010 and related lineages.

**DATA SUMMARY:** Publicly available genomic data were used from three previously published studies (Laycock KM et al., McGee L et al., Khan UB et al.), as well as a set of newly sequenced GBS genomes from clinical strains originating in New York City (NYC). The corresponding accession numbers and detailed information for all strains are provided in the **Table**.

---

Maternal colonization with Group B *Streptococcus* (GBS) with vertical transmission to the neonate is a major source of neonatal morbidity and mortality worldwide.[1, 2] GBS sequence type (ST) 1010 has been identified in studies from several countries, including the United States, Dominican Republic (DR), Netherlands, and Italy.[3, 4] We recently conducted a cross-sectional study of late-pregnancy rectovaginal GBS colonization in the DR and described a local expansion of serotype IV ST1010 GBS, comprising 17% of all isolates in that cohort.[4] Some ST1010 GBS have been noted to contain an *aac*(6′)-*aph*(2″) gene conferring high-level gentamicin resistance (HLGR) carried on a mobile genetic element, though whole genome sequencing data was not available from that study.[3] Here we report an analysis of publicly available and newly sequenced ST1010 isolates and a comparison sample of non-ST1010 GBS focused on the phylogenetic relationship between ST1010 and other GBS lineages, as well as the acquisition of virulence determinants and HLGR.

## METHODS

Laboratory-based surveillance for GBS isolates was conducted under a protocol approved by the institutional review board at NYU Grossman School of Medicine. GBS isolates were collected from the clinical microbiology laboratory and confirmed by growth and phenotype on chromogenic agar (CHROMagar StrepB). DNA extraction was performed using the MagMAX Viral/Pathogen Ultra Nucleic Acid Isolation Kit on a KingFisher Flex automated instrument (ThermoFisher). Library preparation and whole genome sequencing on the NovaSeq 6000 platform were performed at the NYU Genome Technology Core. Adapter removal and quality trimming to a minimum length of 120 bp/read were performed with Trimmomatic 0.36.[5] A complete list of genomes with accession numbers appears in Table 1. We used the SRST2 0.2.0 platform for read mapping to determine multilocus sequence type (MLST) and clonal complex (CC) using allele sequences from pubmlst.org, genomic serotype using the GBS-SBG database, and antibiotic resistance alleles using the database from Metcalf et al.[6-8] Newly identified MLSTs were deposited in the PubMLST database and identifiers were assigned as indicated.[9] SRST2 0.2.0 was also used to detect the presence of *hvgA* and *srr2* alleles using open reading frame sequences from the GBS COH1 genome (NZ_HG939456.1) as well as IS256 using the relevant region of the DR3329 genome (CP129271, bases 197238-180501). IS256 inverted repeat sequences were identified using criteria from Hennig and Ziebuhr.[10]

**Table 1.**
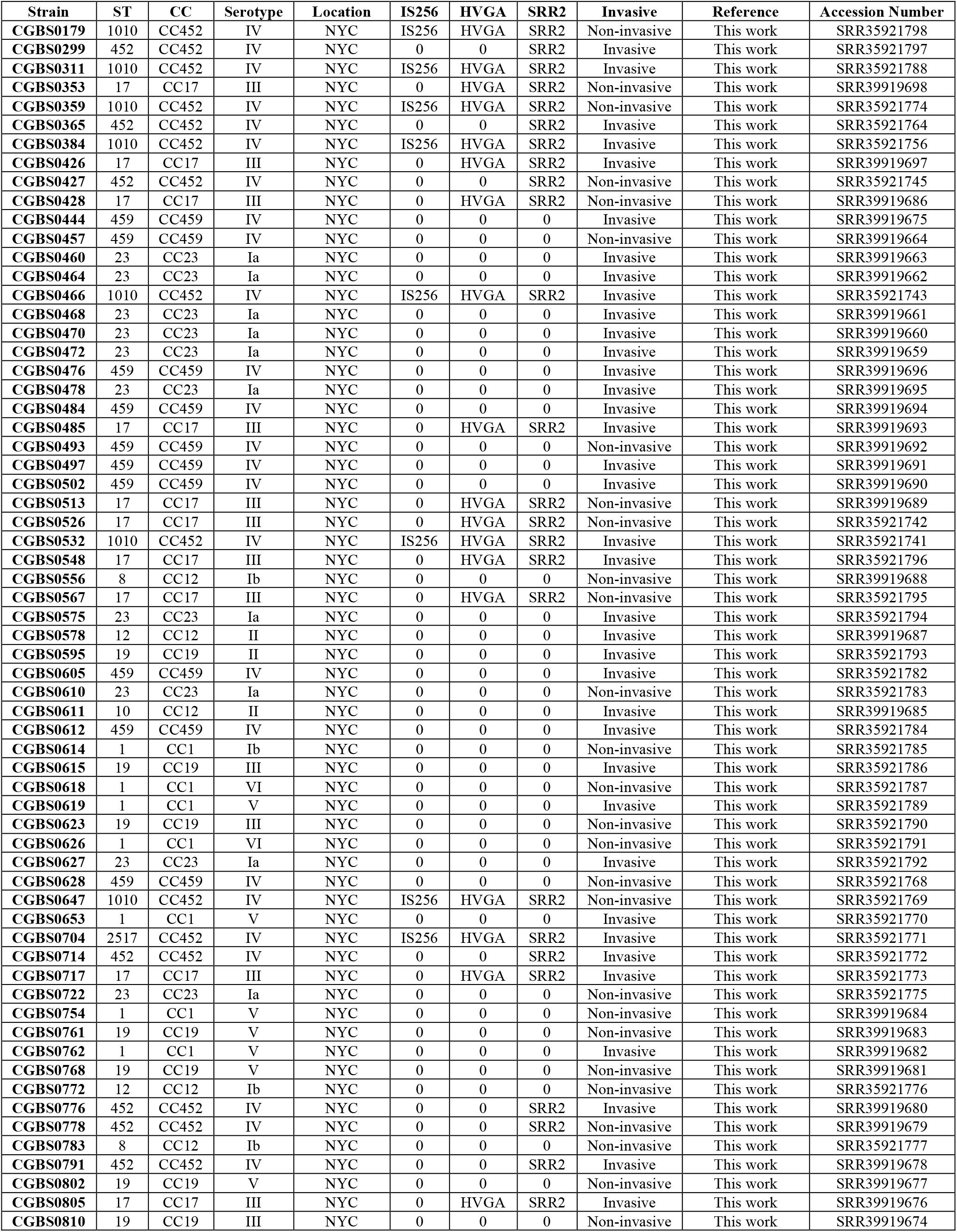
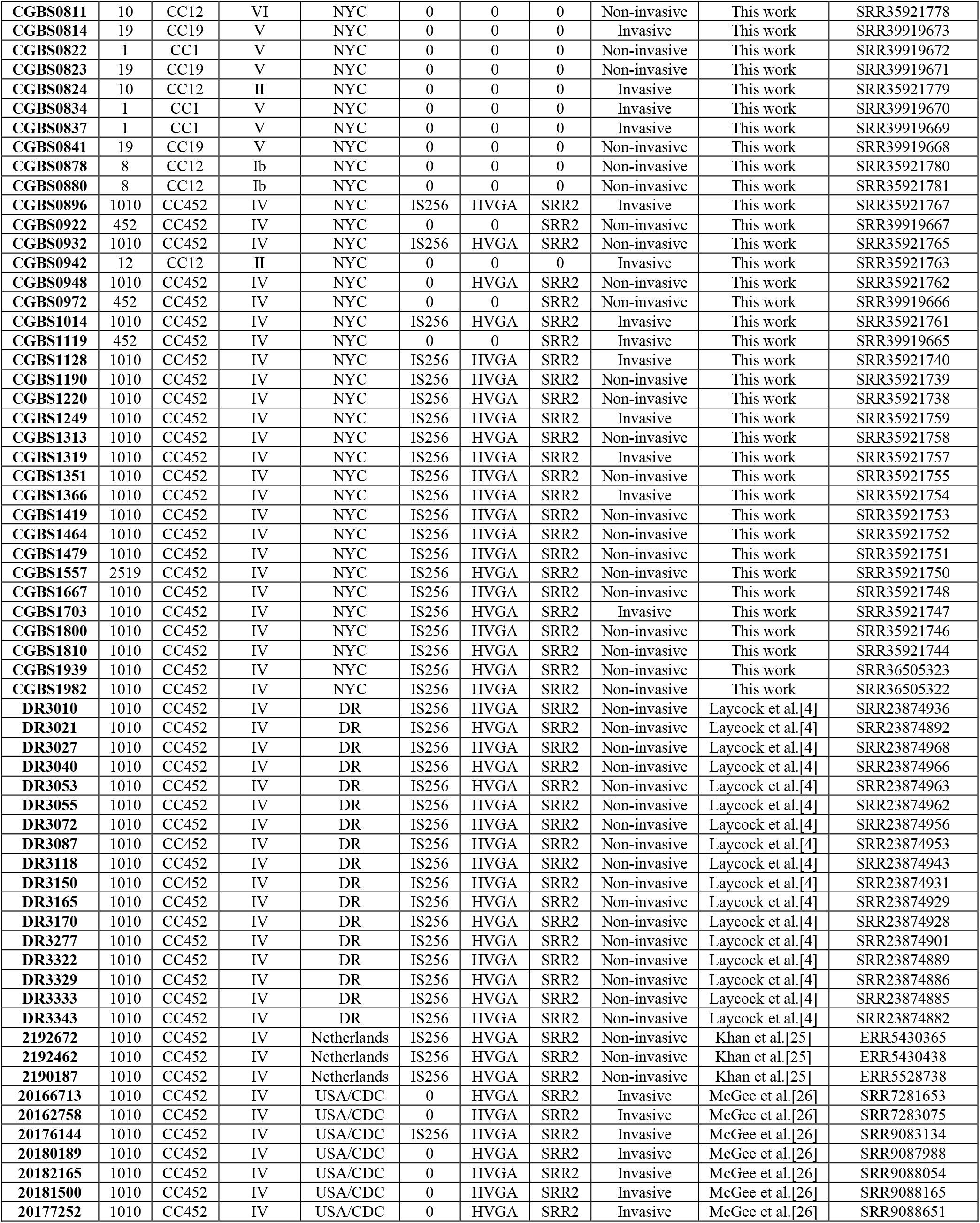
GBS strains used in this study.

| Strain | ST | CC | Serotype | Location | IS256 | HVGA | SRR2 | Invasive | Reference | Accession Number |
| --- | --- | --- | --- | --- | --- | --- | --- | --- | --- | --- |
| CGBS0179 | 1010 | CC452 | IV | NYC | IS256 | HVGA | SRR2 | Non-invasive | This work | SRR35921798 |
| CGBS0299 | 452 | CC452 | IV | NYC | 0 | 0 | SRR2 | Invasive | This work | SRR35921797 |
| CGBS0311 | 1010 | CC452 | IV | NYC | IS256 | HVGA | SRR2 | Invasive | This work | SRR35921788 |
| CGBS0353 | 17 | CC17 | III | NYC | 0 | HVGA | SRR2 | Non-invasive | This work | SRR39919698 |
| CGBS0359 | 1010 | CC452 | IV | NYC | IS256 | HVGA | SRR2 | Non-invasive | This work | SRR35921774 |
| CGBS0365 | 452 | CC452 | IV | NYC | 0 | 0 | SRR2 | Invasive | This work | SRR35921764 |
| CGBS0384 | 1010 | CC452 | IV | NYC | IS256 | HVGA | SRR2 | Invasive | This work | SRR35921756 |
| CGBS0426 | 17 | CC17 | III | NYC | 0 | HVGA | SRR2 | Invasive | This work | SRR39919697 |
| CGBS0427 | 452 | CC452 | IV | NYC | 0 | 0 | SRR2 | Non-invasive | This work | SRR35921745 |
| CGBS0428 | 17 | CC17 | III | NYC | 0 | HVGA | SRR2 | Non-invasive | This work | SRR39919686 |
| CGBS0444 | 459 | CC459 | IV | NYC | 0 | 0 | 0 | Invasive | This work | SRR39919675 |
| CGBS0457 | 459 | CC459 | IV | NYC | 0 | 0 | 0 | Non-invasive | This work | SRR39919664 |
| CGBS0460 | 23 | CC23 | Ia | NYC | 0 | 0 | 0 | Invasive | This work | SRR39919663 |
| CGBS0464 | 23 | CC23 | Ia | NYC | 0 | 0 | 0 | Invasive | This work | SRR39919662 |
| CGBS0466 | 1010 | CC452 | IV | NYC | IS256 | HVGA | SRR2 | Invasive | This work | SRR35921743 |
| CGBS0468 | 23 | CC23 | Ia | NYC | 0 | 0 | 0 | Invasive | This work | SRR39919661 |
| CGBS0470 | 23 | CC23 | Ia | NYC | 0 | 0 | 0 | Invasive | This work | SRR39919660 |
| CGBS0472 | 23 | CC23 | Ia | NYC | 0 | 0 | 0 | Invasive | This work | SRR39919659 |
| CGBS0476 | 459 | CC459 | IV | NYC | 0 | 0 | 0 | Invasive | This work | SRR39919696 |
| CGBS0478 | 23 | CC23 | Ia | NYC | 0 | 0 | 0 | Invasive | This work | SRR39919695 |
| CGBS0484 | 459 | CC459 | IV | NYC | 0 | 0 | 0 | Invasive | This work | SRR39919694 |
| CGBS0485 | 17 | CC17 | III | NYC | 0 | HVGA | SRR2 | Invasive | This work | SRR39919693 |
| CGBS0493 | 459 | CC459 | IV | NYC | 0 | 0 | 0 | Non-invasive | This work | SRR39919692 |
| CGBS0497 | 459 | CC459 | IV | NYC | 0 | 0 | 0 | Invasive | This work | SRR39919691 |
| CGBS0502 | 459 | CC459 | IV | NYC | 0 | 0 | 0 | Invasive | This work | SRR39919690 |
| CGBS0513 | 17 | CC17 | III | NYC | 0 | HVGA | SRR2 | Non-invasive | This work | SRR39919689 |
| CGBS0526 | 17 | CC17 | III | NYC | 0 | HVGA | SRR2 | Non-invasive | This work | SRR35921742 |
| CGBS0532 | 1010 | CC452 | IV | NYC | IS256 | HVGA | SRR2 | Invasive | This work | SRR35921741 |
| CGBS0548 | 17 | CC17 | III | NYC | 0 | HVGA | SRR2 | Invasive | This work | SRR35921796 |
| CGBS0556 | 8 | CC12 | Ib | NYC | 0 | 0 | 0 | Non-invasive | This work | SRR39919688 |
| CGBS0567 | 17 | CC17 | III | NYC | 0 | HVGA | SRR2 | Non-invasive | This work | SRR35921795 |
| CGBS0575 | 23 | CC23 | Ia | NYC | 0 | 0 | 0 | Invasive | This work | SRR35921794 |
| CGBS0578 | 12 | CC12 | II | NYC | 0 | 0 | 0 | Invasive | This work | SRR39919687 |
| CGBS0595 | 19 | CC19 | II | NYC | 0 | 0 | 0 | Invasive | This work | SRR35921793 |
| CGBS0605 | 459 | CC459 | IV | NYC | 0 | 0 | 0 | Invasive | This work | SRR35921782 |
| CGBS0610 | 23 | CC23 | Ia | NYC | 0 | 0 | 0 | Non-invasive | This work | SRR35921783 |
| CGBS0611 | 10 | CC12 | II | NYC | 0 | 0 | 0 | Invasive | This work | SRR39919685 |
| CGBS0612 | 459 | CC459 | IV | NYC | 0 | 0 | 0 | Invasive | This work | SRR35921784 |
| CGBS0614 | 1 | CC1 | Ib | NYC | 0 | 0 | 0 | Non-invasive | This work | SRR35921785 |
| CGBS0615 | 19 | CC19 | III | NYC | 0 | 0 | 0 | Invasive | This work | SRR35921786 |
| CGBS0618 | 1 | CC1 | VI | NYC | 0 | 0 | 0 | Non-invasive | This work | SRR35921787 |
| CGBS0619 | 1 | CC1 | V | NYC | 0 | 0 | 0 | Invasive | This work | SRR35921789 |
| CGBS0623 | 19 | CC19 | III | NYC | 0 | 0 | 0 | Non-invasive | This work | SRR35921790 |
| CGBS0626 | 1 | CC1 | VI | NYC | 0 | 0 | 0 | Non-invasive | This work | SRR35921791 |
| CGBS0627 | 23 | CC23 | Ia | NYC | 0 | 0 | 0 | Invasive | This work | SRR35921792 |
| CGBS0628 | 459 | CC459 | IV | NYC | 0 | 0 | 0 | Non-invasive | This work | SRR35921768 |
| CGBS0647 | 1010 | CC452 | IV | NYC | IS256 | HVGA | SRR2 | Non-invasive | This work | SRR35921769 |
| CGBS0653 | 1 | CC1 | V | NYC | 0 | 0 | 0 | Invasive | This work | SRR35921770 |
| CGBS0704 | 2517 | CC452 | IV | NYC | IS256 | HVGA | SRR2 | Invasive | This work | SRR35921771 |
| CGBS0714 | 452 | CC452 | IV | NYC | 0 | 0 | SRR2 | Invasive | This work | SRR35921772 |
| CGBS0717 | 17 | CC17 | III | NYC | 0 | HVGA | SRR2 | Invasive | This work | SRR35921773 |
| CGBS0722 | 23 | CC23 | Ia | NYC | 0 | 0 | 0 | Non-invasive | This work | SRR35921775 |
| CGBS0754 | 1 | CC1 | V | NYC | 0 | 0 | 0 | Non-invasive | This work | SRR39919684 |
| CGBS0761 | 19 | CC19 | V | NYC | 0 | 0 | 0 | Non-invasive | This work | SRR39919683 |
| CGBS0762 | 1 | CC1 | V | NYC | 0 | 0 | 0 | Invasive | This work | SRR39919682 |
| CGBS0768 | 19 | CC19 | V | NYC | 0 | 0 | 0 | Non-invasive | This work | SRR39919681 |
| CGBS0772 | 12 | CC12 | Ib | NYC | 0 | 0 | 0 | Non-invasive | This work | SRR35921776 |
| CGBS0776 | 452 | CC452 | IV | NYC | 0 | 0 | SRR2 | Invasive | This work | SRR39919680 |
| CGBS0778 | 452 | CC452 | IV | NYC | 0 | 0 | SRR2 | Non-invasive | This work | SRR39919679 |
| CGBS0783 | 8 | CC12 | Ib | NYC | 0 | 0 | 0 | Non-invasive | This work | SRR35921777 |
| CGBS0791 | 452 | CC452 | IV | NYC | 0 | 0 | SRR2 | Invasive | This work | SRR39919678 |
| CGBS0802 | 19 | CC19 | V | NYC | 0 | 0 | 0 | Non-invasive | This work | SRR39919677 |
| CGBS0805 | 17 | CC17 | III | NYC | 0 | HVGA | SRR2 | Invasive | This work | SRR39919676 |
| CGBS0810 | 19 | CC19 | III | NYC | 0 | 0 | 0 | Non-invasive | This work | SRR39919674 |
| CGBS0811 | 10 | CC12 | VI | NYC | 0 | 0 | 0 | Non-invasive | This work | SRR35921778 |
| CGBS0814 | 19 | CC19 | V | NYC | 0 | 0 | 0 | Invasive | This work | SRR39919673 |
| CGBS0822 | 1 | CC1 | V | NYC | 0 | 0 | 0 | Non-invasive | This work | SRR39919672 |
| CGBS0823 | 19 | CC19 | V | NYC | 0 | 0 | 0 | Non-invasive | This work | SRR39919671 |
| CGBS0824 | 10 | CC12 | II | NYC | 0 | 0 | 0 | Invasive | This work | SRR35921779 |
| CGBS0834 | 1 | CC1 | V | NYC | 0 | 0 | 0 | Invasive | This work | SRR39919670 |
| CGBS0837 | 1 | CC1 | V | NYC | 0 | 0 | 0 | Invasive | This work | SRR39919669 |
| CGBS0841 | 19 | CC19 | V | NYC | 0 | 0 | 0 | Non-invasive | This work | SRR39919668 |
| CGBS0878 | 8 | CC12 | Ib | NYC | 0 | 0 | 0 | Non-invasive | This work | SRR35921780 |
| CGBS0880 | 8 | CC12 | Ib | NYC | 0 | 0 | 0 | Non-invasive | This work | SRR35921781 |
| CGBS0896 | 1010 | CC452 | IV | NYC | IS256 | HVGA | SRR2 | Invasive | This work | SRR35921767 |
| CGBS0922 | 452 | CC452 | IV | NYC | 0 | 0 | SRR2 | Non-invasive | This work | SRR39919667 |
| CGBS0932 | 1010 | CC452 | IV | NYC | IS256 | HVGA | SRR2 | Non-invasive | This work | SRR35921765 |
| CGBS0942 | 12 | CC12 | II | NYC | 0 | 0 | 0 | Invasive | This work | SRR35921763 |
| CGBS0948 | 1010 | CC452 | IV | NYC | 0 | HVGA | SRR2 | Non-invasive | This work | SRR35921762 |
| CGBS0972 | 452 | CC452 | IV | NYC | 0 | 0 | SRR2 | Non-invasive | This work | SRR39919666 |
| CGBS1014 | 1010 | CC452 | IV | NYC | IS256 | HVGA | SRR2 | Invasive | This work | SRR35921761 |
| CGBS1119 | 452 | CC452 | IV | NYC | 0 | 0 | SRR2 | Invasive | This work | SRR39919665 |
| CGBS1128 | 1010 | CC452 | IV | NYC | IS256 | HVGA | SRR2 | Invasive | This work | SRR35921740 |
| CGBS1190 | 1010 | CC452 | IV | NYC | IS256 | HVGA | SRR2 | Non-invasive | This work | SRR35921739 |
| CGBS1220 | 1010 | CC452 | IV | NYC | IS256 | HVGA | SRR2 | Non-invasive | This work | SRR35921738 |
| CGBS1249 | 1010 | CC452 | IV | NYC | IS256 | HVGA | SRR2 | Invasive | This work | SRR35921759 |
| CGBS1313 | 1010 | CC452 | IV | NYC | IS256 | HVGA | SRR2 | Non-invasive | This work | SRR35921758 |
| CGBS1319 | 1010 | CC452 | IV | NYC | IS256 | HVGA | SRR2 | Invasive | This work | SRR35921757 |
| CGBS1351 | 1010 | CC452 | IV | NYC | IS256 | HVGA | SRR2 | Non-invasive | This work | SRR35921755 |
| CGBS1366 | 1010 | CC452 | IV | NYC | IS256 | HVGA | SRR2 | Invasive | This work | SRR35921754 |
| CGBS1419 | 1010 | CC452 | IV | NYC | IS256 | HVGA | SRR2 | Non-invasive | This work | SRR35921753 |
| CGBS1464 | 1010 | CC452 | IV | NYC | IS256 | HVGA | SRR2 | Non-invasive | This work | SRR35921752 |
| CGBS1479 | 1010 | CC452 | IV | NYC | IS256 | HVGA | SRR2 | Non-invasive | This work | SRR35921751 |
| CGBS1557 | 2519 | CC452 | IV | NYC | IS256 | HVGA | SRR2 | Non-invasive | This work | SRR35921750 |
| CGBS1667 | 1010 | CC452 | IV | NYC | IS256 | HVGA | SRR2 | Non-invasive | This work | SRR35921748 |
| CGBS1703 | 1010 | CC452 | IV | NYC | IS256 | HVGA | SRR2 | Invasive | This work | SRR35921747 |
| CGBS1800 | 1010 | CC452 | IV | NYC | IS256 | HVGA | SRR2 | Non-invasive | This work | SRR35921746 |
| CGBS1810 | 1010 | CC452 | IV | NYC | IS256 | HVGA | SRR2 | Non-invasive | This work | SRR35921744 |
| CGBS1939 | 1010 | CC452 | IV | NYC | IS256 | HVGA | SRR2 | Non-invasive | This work | SRR36505323 |
| CGBS1982 | 1010 | CC452 | IV | NYC | IS256 | HVGA | SRR2 | Non-invasive | This work | SRR36505322 |
| DR3010 | 1010 | CC452 | IV | DR | IS256 | HVGA | SRR2 | Non-invasive | Laycock et al.[4] | SRR23874936 |
| DR3021 | 1010 | CC452 | IV | DR | IS256 | HVGA | SRR2 | Non-invasive | Laycock et al.[4] | SRR23874892 |
| DR3027 | 1010 | CC452 | IV | DR | IS256 | HVGA | SRR2 | Non-invasive | Laycock et al.[4] | SRR23874968 |
| DR3040 | 1010 | CC452 | IV | DR | IS256 | HVGA | SRR2 | Non-invasive | Laycock et al.[4] | SRR23874966 |
| DR3053 | 1010 | CC452 | IV | DR | IS256 | HVGA | SRR2 | Non-invasive | Laycock et al.[4] | SRR23874963 |
| DR3055 | 1010 | CC452 | IV | DR | IS256 | HVGA | SRR2 | Non-invasive | Laycock et al.[4] | SRR23874962 |
| DR3072 | 1010 | CC452 | IV | DR | IS256 | HVGA | SRR2 | Non-invasive | Laycock et al.[4] | SRR23874956 |
| DR3087 | 1010 | CC452 | IV | DR | IS256 | HVGA | SRR2 | Non-invasive | Laycock et al.[4] | SRR23874953 |
| DR3118 | 1010 | CC452 | IV | DR | IS256 | HVGA | SRR2 | Non-invasive | Laycock et al.[4] | SRR23874943 |
| DR3150 | 1010 | CC452 | IV | DR | IS256 | HVGA | SRR2 | Non-invasive | Laycock et al.[4] | SRR23874931 |
| DR3165 | 1010 | CC452 | IV | DR | IS256 | HVGA | SRR2 | Non-invasive | Laycock et al.[4] | SRR23874929 |
| DR3170 | 1010 | CC452 | IV | DR | IS256 | HVGA | SRR2 | Non-invasive | Laycock et al.[4] | SRR23874928 |
| DR3277 | 1010 | CC452 | IV | DR | IS256 | HVGA | SRR2 | Non-invasive | Laycock et al.[4] | SRR23874901 |
| DR3322 | 1010 | CC452 | IV | DR | IS256 | HVGA | SRR2 | Non-invasive | Laycock et al.[4] | SRR23874889 |
| DR3329 | 1010 | CC452 | IV | DR | IS256 | HVGA | SRR2 | Non-invasive | Laycock et al.[4] | SRR23874886 |
| DR3333 | 1010 | CC452 | IV | DR | IS256 | HVGA | SRR2 | Non-invasive | Laycock et al.[4] | SRR23874885 |
| DR3343 | 1010 | CC452 | IV | DR | IS256 | HVGA | SRR2 | Non-invasive | Laycock et al.[4] | SRR23874882 |
| 2192672 | 1010 | CC452 | IV | Netherlands | IS256 | HVGA | SRR2 | Non-invasive | Khan et al.[25] | ERR5430365 |
| 2192462 | 1010 | CC452 | IV | Netherlands | IS256 | HVGA | SRR2 | Non-invasive | Khan et al.[25] | ERR5430438 |
| 2190187 | 1010 | CC452 | IV | Netherlands | IS256 | HVGA | SRR2 | Non-invasive | Khan et al.[25] | ERR5528738 |
| 20166713 | 1010 | CC452 | IV | USA/CDC | 0 | HVGA | SRR2 | Invasive | McGee et al.[26] | SRR7281653 |
| 20162758 | 1010 | CC452 | IV | USA/CDC | 0 | HVGA | SRR2 | Invasive | McGee et al.[26] | SRR7283075 |
| 20176144 | 1010 | CC452 | IV | USA/CDC | IS256 | HVGA | SRR2 | Invasive | McGee et al.[26] | SRR9083134 |
| 20180189 | 1010 | CC452 | IV | USA/CDC | 0 | HVGA | SRR2 | Invasive | McGee et al.[26] | SRR9087988 |
| 20182165 | 1010 | CC452 | IV | USA/CDC | 0 | HVGA | SRR2 | Invasive | McGee et al.[26] | SRR9088054 |
| 20181500 | 1010 | CC452 | IV | USA/CDC | 0 | HVGA | SRR2 | Invasive | McGee et al.[26] | SRR9088165 |
| 20177252 | 1010 | CC452 | IV | USA/CDC | 0 | HVGA | SRR2 | Invasive | McGee et al.[26] | SRR9088651 |

Short-read sequences were assembled using AbySS 2.1.1, assessed for quality with QUAST 5.0.2, and annotated with prokka 1.14.6 using GBS 2603V/R (NC_004116) as a reference sequence.[11-13] A core genome alignment was generated with roary 3.12.0, and a core phylogeny with IQ-TREE 3.0.1 using modelfinder (best fit model GTR+F+I+R5) and visualized on the Microreact platform.[14]

A whole-genome, kmer-based alignment of members of the ST1010 clade (including single-locus variants) was generated using SKA2.[15] Regions of recombination within the ST1010 clade were identified using Gubbins 3.3.5, and a recombination-masked tree was generated with RAXML-MG running in the Gubbins package.[16, 17] The closed DR3329 genome (CP129271) was used as the reference. Visualization was performed with Phandango 1.3.1.[18]

## RESULTS

The dataset included 57 invasive disease isolates and 70 maternal carriage (non-invasive) isolates (**Table**). Two of the newly sequenced isolates collected in NYC were identified as single locus variants of ST1010 at loci *adhP* (5→461) and *glnA* (2→9). These were submitted to PubMLST and assigned STs 2517 and 2519 respectively.

**Fig. 1a** shows the core phylogeny of the set of GBS isolates, as well as the presence of key virulence determinant genes *hvgA* and *srr2* and the IS256 mobile element conferring HLGR in each isolate. Consistent with prior studies, ST1010 appears closely related to ST17 and ST452. All ST1010 and ST17 isolates grouped into a common clade, branching distal to ST452 (**Fig. 1b**). All ST1010 and ST17 isolates possess both *hvgA* and *srr2*. The *srr2* gene is also present in all ST452 isolates. The IS256 mobile element was present in most (48/55, 87%) ST1010 isolates, along with the two novel single locus variants of ST1010 (ST2517 and ST2519). The IS256-negative ST1010 isolates formed a clade distinct from the IS256-containing strains (**Fig. 1c**).

**Figure 1.**
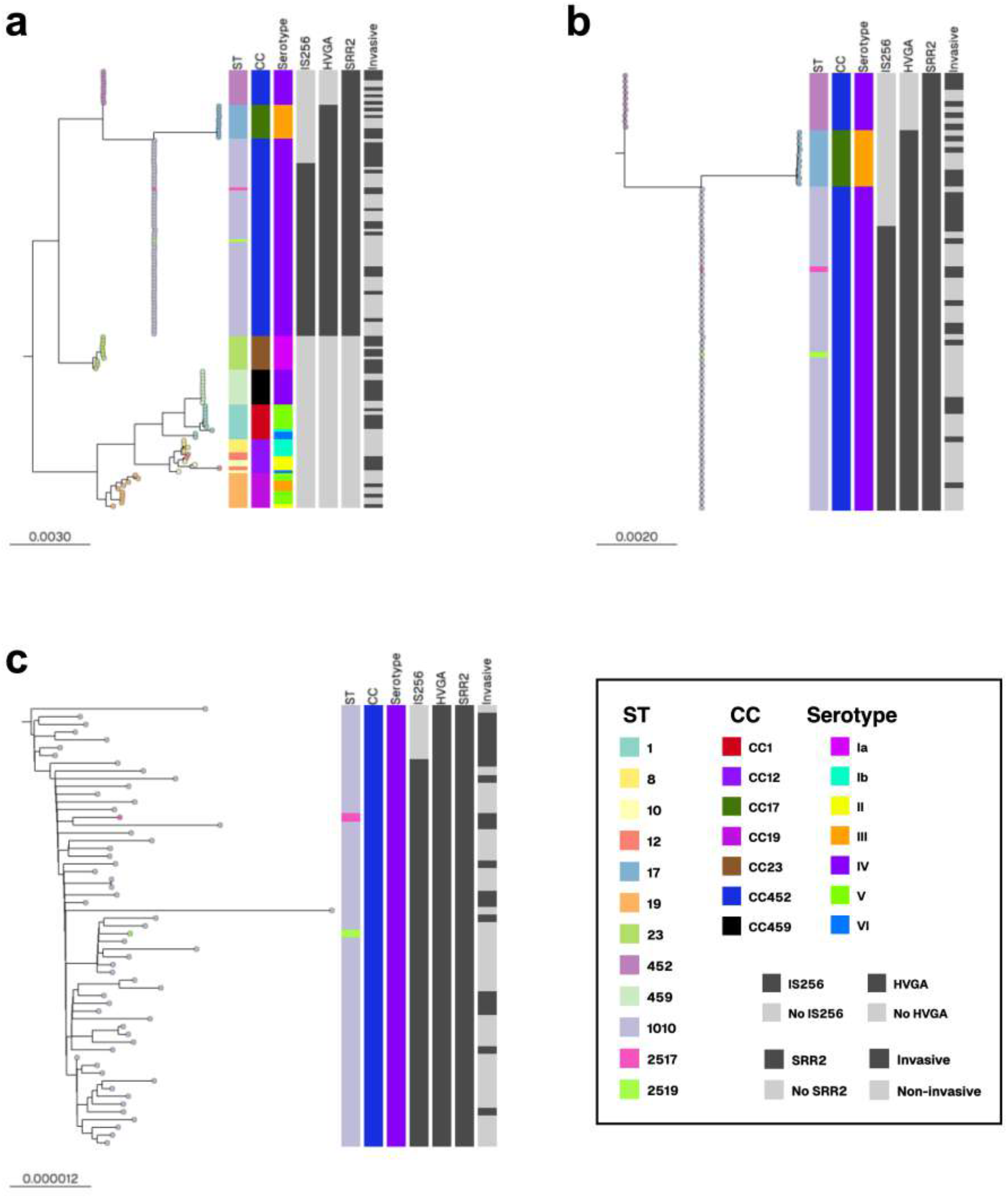
(**a**) Core genome tree of GBS isolates with associated metadata. (**b**) Subtree of ST1010, ST452, and ST17 demonstrates that ST1010 and ST17 form a clade that shares a common ancestor with ST452. (**c**) Subtree of ST1010 isolates reveals that isolates with and without IS256 cluster into separate clades.

We visualized predicted recombination events in the ST1010/ST2517/ST2519 group using Gubbins (**Fig. 2a**). Limited areas of recombination were identified, corresponding to genomic regions enriched in predicted mobile genetic element components. The clade containing IS256-negative ST1010 isolates harbors a unique region of recombination within the large integrative and conjugative element ICESag9931 identified by Creti et al.[3] ICESag9931 contains the *aac*(6′)-*aph*(2″) gene that confers HLGR within a predicted IS256 insertion sequence (**Fig. 2b**). Together, these data suggest acquisition and expansion of a distinct GBS ST1010 lineage with the genetic machinery for HLGR as well as multiple virulence determinants not previously appreciated outside of CC17 GBS.

**Figure 2.**
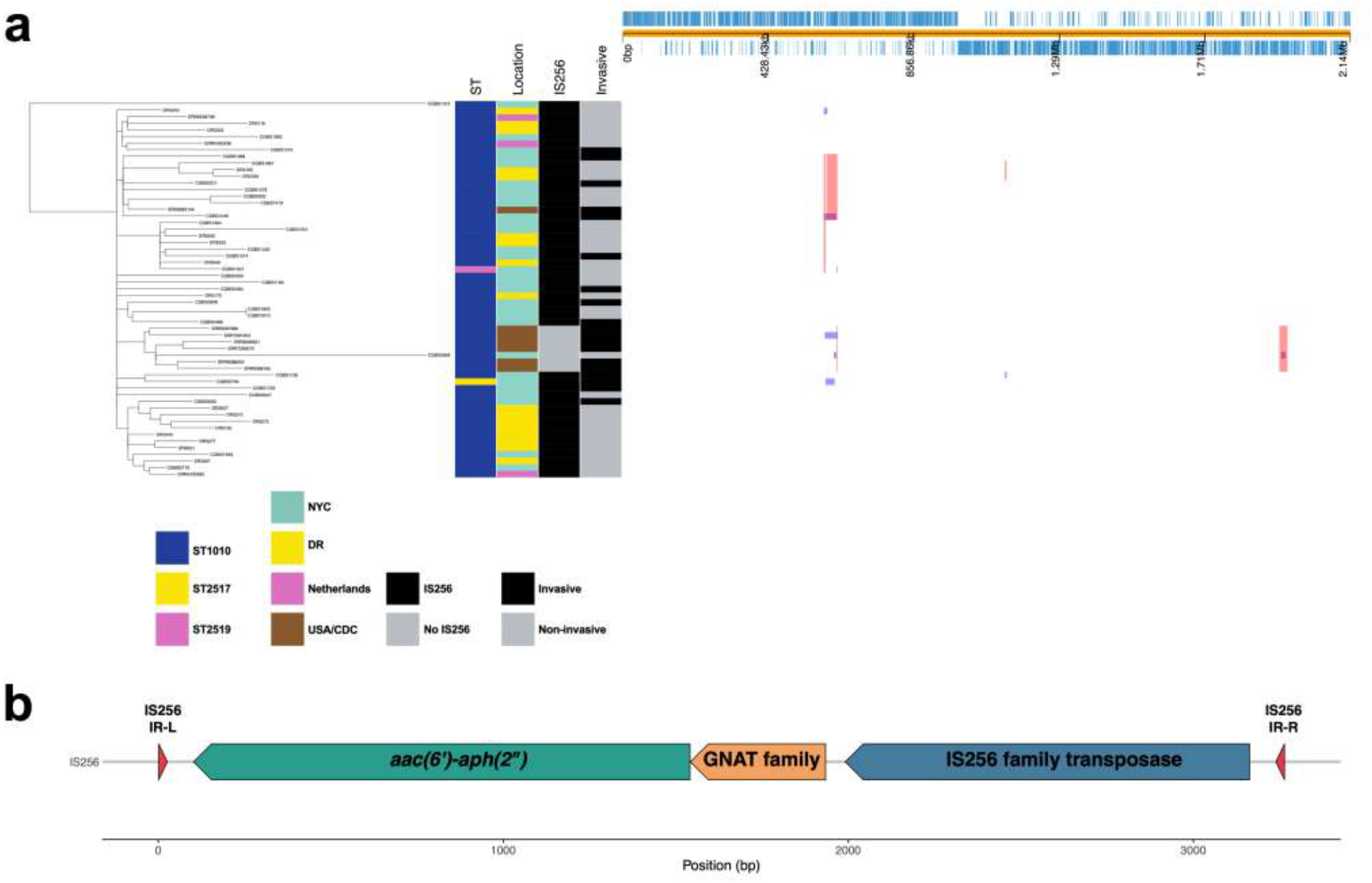
(**a**) Identification of putative areas of recombination within the clade containing ST1010 and its newly identified single-locus variants. Sites of recombination were identified using Gubbins. A recombination-masked tree was constructed (at left) and displayed with metadata (columns at middle) and areas of recombination mapped to the linearized DR3329 genome (at top). Recombination areas were in regions annotated as mobile genetic elements, including ICESag9931 (far right). (**b**) Map of the IS256 element in the DR3329 (ST1010) genome containing consensus IS256 inverted repeats at left (IS256 IR-L) and right (IS256IR-R), an *aac(6’)-aph(2”)* gene conferring HLGR, a GNAT family acetyltransferase, and an IS256 transposase as indicated.

## DISCUSSION

This study describes the phylogenetic and genomic characteristics of a diverse set of GBS ST1010 isolates and introduces two novel sequence types (ST2517 and ST2519) identified as single locus variants of ST1010. The results of our core phylogeny show ST1010 and ST17 in a common clade following divergence from their shared common ancestor with ST452. Genes encoding key virulence determinants, the HvgA adhesin and the serine-rich repeat protein Srr2, were present in all isolates in this common clade, while *hvgA* was absent in ST452 isolates, suggesting *hvgA* acquisition prior to divergence of ST1010 and ST17. HvgA and Srr2 are surface proteins that promote adhesion and internalization of CC17 strains into brain endothelial cells, facilitating hypervirulence and associated with poorer clinical outcomes.[19, 20] ST17 GBS is especially prevalent in cases of neonatal meningitis in both early-onset and late-onset GBS disease and is considered a “hypervirulent” lineage.[21-23] This study demonstrates that ST1010 is not only closely related to ST17 but also shares key virulence factors and therefore may possess similar virulence phenotypes.

Both ST452 and ST1010 express a serotype IV polysaccharide capsule, while ST17 possesses the serotype III capsule. A prior study describing ST452 suggested that ST452 is a capsule switch strain arising from a large recombination event between CC23 and CC17 lineages.[24] However, the relationship between ST452, ST1010, and ST17 strains described above suggests instead that ST17 may have arisen from a common ancestor of serotype IV lineages ST452 and ST1010 and subsequently acquired the serotype III capsule machinery.

In addition, our results suggest that following the divergence of the ST1010 group, acquisition of a mobile genetic element containing an *aac*(6′)-*aph*(2″) gene likely led to a HLGR phenotype. Within ST1010, strains containing this element are numerically dominant, suggesting that most ST1010 strains that are emerging internationally carry a concerning combination of antibiotic resistance and virulence determinants. These results have implications for ongoing GBS surveillance and understanding emergence of novel GBS lineages.

## Abbreviations

CC: clonal complex
CDC: Centers for Disease Control and Prevention
DR: Dominican Republic
GBS: Group B *Streptococcus*
HLGR: high-level gentamicin resistance
IS: insertion sequence
ML: maximum likelihood
MLST: multilocus sequence type
NYC: New York City
ST: sequence type

## ACKNOWLEDGEMENTS

We acknowledge the NYU Langone Antimicrobial-Resistant Pathogens Program (Dr. Bo Shopsin) and the NYU Langone Health Genome Technology Center for assistance with GBS whole-genome sequencing.

## AUTHOR CONTRIBUTIONS

ACF: conceptualization, funding acquisition, data curation, formal analysis, investigation, methodology, validation, visualization, writing – original draft, writing – review & editing

SH: data curation, formal analysis, investigation, methodology, validation, visualization, writing – review & editing

CO: investigation, methodology, resources, writing – review & editing AD: investigation, methodology, visualization, writing – review & editing

HT: conceptualization, data curation, formal analysis, investigation, methodology, project administration, resources, validation, visualization, writing – review & editing

AJR: conceptualization, funding acquisition, data curation, formal analysis, investigation, methodology, project administration, resources, validation, visualization, writing – original draft, writing – review & editing

